# Exploring Prime Editing strategies for genetic correction of a Ceroid Neuronal Lipofuscinosis type 2 disease causing variant

**DOI:** 10.1101/2025.10.02.680174

**Authors:** Heshadi Primrose Mandalawatta, KC Rajendra, Kirsten A Fairfax, Alex W Hewitt

**Affiliations:** Menzies Institute for Medical Research, University of Tasmania, Australia; School of Medicine, University of Tasmania, Australia

## Abstract

Neuronal ceroid lipofuscinosis type 2 (CLN2) is a rare, fatal paediatric neurodegenerative genetic disease that causes progressive psychomotor decline, epilepsy, speech impairment, vision loss, and premature death by late childhood or early adolescence. It is caused by mutations in the *TPP1* gene, which encodes the tripeptidyl peptidase 1 enzyme.

To date, enzyme replacement therapy (ERT) with recombinant human TPP1 is the only approved treatment available and it does not appear to have a curative effect on disease progression or rescuing complex motor functions. This underscores the importance of developing more effective treatment strategies for CLN2 disease. In the quest for an alternative therapy, we here seek to utilise CRISPR based prime editing (PE) to correct one of the most frequent TPP1 mutations of c.622 C>T (p. Arg208Ter).

In cultured mammalian HEK293a cells we screened 16 different PE and pegRNA combinations, achieving on-target editing efficiencies up to 27.36 %. To enable efficient *in vivo* gene editing, we packaged the most promising PE and pegRNA combination using engineered virus-like particles (eVLPs) which resulted in 12.6 % on-target editing. Given the robust on-target editing and reduced risk of bystander editing, PE is deemed suitable for *in vivo* testing in the pre-clinical models for CLN2 disease.

Overall, our findings establish the proof of concept for CRISPR-based prime editing to correct pathogenic human *TPP1* nonsense mutation of c.622 C>T (p. R208X) and provide a basis for further investigations of PE as a genetic treatment for CLN2 disease.

## MAIN

The neuronal ceroid lipofuscinoses (NCLs), also known as Batten disease, are a group of inherited, lysosomal storage disorders (LSDs) that are profoundly neurodegenerative. To-date 14 genetically distinct NCL types have been identified. These subtypes share similar symptoms such as recurrent seizures, visual impairment, cognitive and motor decline and ultimately lead to premature death ^1,2^. Classic late infantile NCL, or CLN2 disease, is one of the most common forms of NCLs which is caused by the deficiency of tripeptidyl peptidase 1 (TPP1) enzyme. The splice mutation of c.509-1G>C and c.622 C>T (p.Arg208Ter) are the most prevalent forms of CLN2 disease-causing mutations ^3–5^. These mutations affect endo-lysosomal function and result in progressive accumulation of intracellular auto-fluorescent storage material. The composition and ultrastructural profiles of these auto-fluorescent storage materials are varied among the NCL subtypes ^6,7^. While there is currently no cure for CLN2 disease, early intervention of the Enzyme replacement therapy (ERT) of the human TPP1 proenzyme was shown to alleviate disease progression ^8^. This highlights the importance of developing alternative therapeutic approaches to treat the CLN2 disease. Here, we describe the application of PE to correct the murine mutation (p.R207X) equivalent to the pathogenic human *TPP1* c.622 C>T (p. R208X) mutation.

Prime editor is a versatile editing agent that can perform a diverse and precise editing of any of the 12 possible single nucleotide conversions, substitutions, insertions and deletions without inducing double stranded DNA breaks ^9^. Originally, the prime editor 1 (PE1) was developed fusing a catalytically inactive *Streptococcus pyogenes* Cas9 nickase mutant (H840A) to an engineered reverse transcriptase (RT) derived from Moloney murine leukaemia virus (M-MLV). Similar to a single guide RNA, a prime editing guide RNA (pegRNA) navigates the PE complex to the target genomic location. The 3’ extension of the pegRNA contains a reverse transcription template region which provides instruction for the desired edit and a primer binding site (PBS) complementary to the target sequence. To initiate prime editing nCas9 introduces a single nick near the PAM site on the sense DNA strand and the RT utilises the PBS sequence of the pegRNA as a template to extend the 3′**-**end of the nicked DNA strand, introducing the desired edits into the new 3′ flap. The cellular DNA replication and repair pathways preferentially retain the 3’ flap over the existing 5’ flap. Finally, the mismatch repair mechanism corrects the unedited strand.^9^

Our results demonstrate that PE can precisely correct the equivalent murine mutation (p.R207X) of the pathogenic human *TPP1* nonsense mutation of c.622 C>T (p. R208X) *in vitro* without detectable bystander edits.

## RESULTS

When designing a prime editing experiment, careful evaluation of the pegRNA design, structure and target scope and formation of silent mutations contribute to success of the experiment. Though not all these aspects allow for straightforward decisions, structural optimisations are particularly helpful to expand the editing scope.^10^ For example, when selecting an appropriate PE and pegRNA combination, the nature of the target site, chromatin accessibility,^11^ protospacer adjacent motif (PAM) to edit distance (if using Cas9 variants with NG/NGG PAMs) ^12^ and propensity of indel formation are key determinants.

### Initial screening

Our goal was to evaluate PE as a potential transformative therapy for CLN2 batten disease. Since the initial report, PE has been progressively developed for correction of a vast array of disease-causing mutations that were previously inaccessible. Among the reported PE1-PE7 variants,^9,13–15^ the PE2 variant, that encodes a P2A-blasticidin resistance marker (BSR) fusion under a cytomegalovirus promoter was selected over other systems as it offers the benefit of using a single pegRNA.^9^

To investigate the PE-mediated correction of the mutation, cultured HEK293A cells (harbouring homozygous p.R207X mutation) were transfected with plasmids expressing the PE2 and the pegRNA and genomic DNA was harvested 96 hours post-transfection. Although the initial screening of PE2-SpCas9 and unmodified pegRNA resulted in a low level of editing (0.66 %, s.d = 0.577) of editing, we continued to focus on incorporating other means of PE advancements such as structural modifications of the PE and pegRNA.^10,16^

### Structural optimisation of pegRNA

In the prime editing mechanism, the pegRNA plays a vital role: navigating the editor to the target sequence and encoding the desired edit. Therefore, selecting a suitable pegRNA architecture necessitates thorough and careful optimisation. Indeed, we have synergistically optimized both PE and the pegRNA to maximise the editing versatility. As such the pegRNA modifications included integration of structured RNA motifs into the 3′end of pegRNAs^17^ and fusion of RNA-binding protein La to protect the end of the pegRNA.^15^ Also, to assess the impact of modifying the primer binding site (PBS) and reverse transcriptase template (RTT) of the pegRNA we designed an oligopool of different PBS and RTT lengths^18^ and also introduced same-sense mutations at proper positions in the reverse-transcription template of pegRNA.^19^ Additionally, we utilised the dual pegRNA strategy and also designed pegRNAs with different edit-to-nick distances.^20^

### Engineered pegRNA

The binding stability of the 3’ end of the pegRNA to the target sequence is critical for the PE editing efficiency and degradation of the 3’ end of pegRNA may potentially impede prime editing. Therefore, to circumvent this, Nelson *et al* (2022) introduced a structured RNA motif (trimmed evopreQ1 or tevopreQ1) at the 3’ of the PBS which was shown to stabilise the pegRNA structure and subsequently prevent it from cellular exonuclease degradation. These engineered pegRNAs (epegRNA) with the 3’ motif have been observed to improve prime editing efficiency over the unmodified pegRNA.^17^

Previous studies have reported improved *in vitro* editing for epegRNA across multiple loci and cell types,^10^ and in this study the *in vitro* evaluation of PE2-SpCas9 with epegRNA yielded up to 5.66 % (s.d = 1.15) editing. Given that epegRNA resulted in an 8.6-fold increase in prime editing compared to unmodified pegRNA (control), we speculated that further structural optimisation of PE components could improve prime editing efficiency.

### PAM flexible Cas9 variants

The Cas9 protein derived from the bacteria *Streptococcus pyogenes* (SpCas9) has a relatively short canonical PAM recognition site 5′-NGG-3′, where N could be any nucleotide.^21^ However, the strict NGG PAM preference often limits the target scope of the PE, thereby excluding many disease relevant targets from being candidates for prime editing. In contrast, engineered SpCas9 variants with relaxed PAM preferences such as SpG-Cas9 (5’-NGN-3’ PAM)^22^ and SpRY-Cas9 (5’-NYN-3’ where Y is C or T) recognise minimal PAM sequences (NG) or function independently of the PAM.^23^ Here, the Cas9 module of the PE2 was substituted with PAM-flexible Cas9 variants SpG and SpRY and their editing efficiencies were compared side-by-side.

Interestingly we observed improved and measurable levels of editing compared to PE2-SpCas9 (control). On average, PE2-SpRY-Cas9 prime editing with unmodified pegRNA resulted in 7.23 % (s.d = 0.28, p-value < 0.0001 by two-way ANOVA) and PE2-SpG-Cas9 outperformed PE2-SpRY-Cas9 by 1.2-fold (8.71 %, s.d = 0.63, p-value < 0.0001 by two-way ANOVA) (**Figure 1a**). These results highlight the key functional differences among different Cas9 variants and such differences in genome editing arise from the differences between the target specificity and PAM compatibility.^24^ Moreover, 3′ extension of the pegRNA has shown to reduce SpCas9’s efficiency in forming the R-loop or performing cleavage suggesting that SpRY-Cas9 may have better compatibility with pegRNA.^25^ This further highlights the importance of Cas9-pegRNA complexing, DNA target engagement or DNA nicking kinetics to improve the magnitude of PE efficiency.^12^

**Figure 1.**
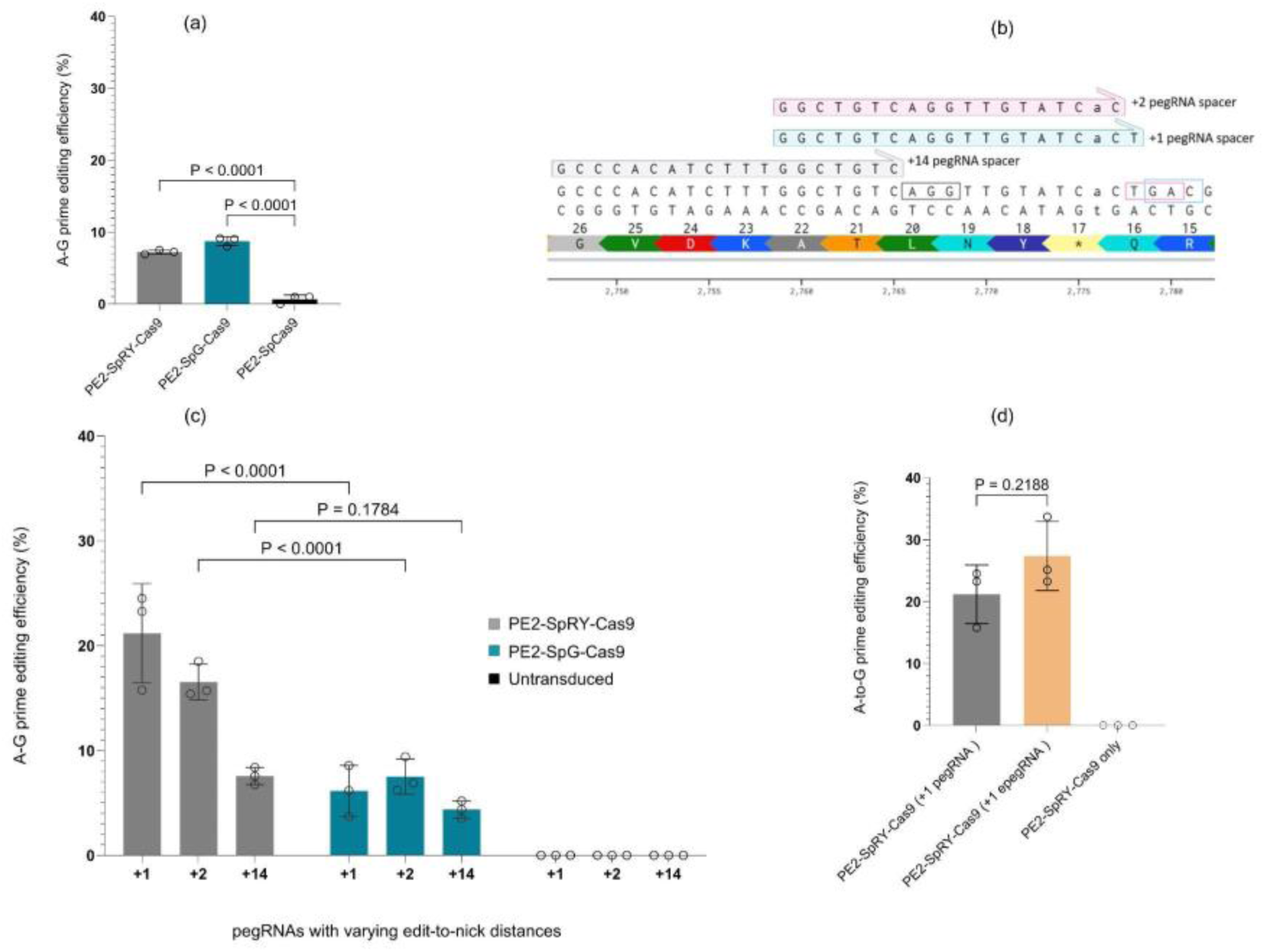
Comparison of in vitro A-to-G editing efficiencies of PE2-SpG-Cas9 and PE2-SpRY-Cas9 when paired with unmodified pegRNA combinations. a. Bar graph showing mean A-to-G on-target editing of PE2-SpG-Cas9 and PE2-SpRY-Cas9 compared to that of PE2-SpCas9 (0.66 %, s.d = 0.57, control). PE2-SpRY-Cas9 and PE2-SpG-Cas9 resulted in and 7.23 % (s.d = 0.28, ****p-value < 0.0001) and 8.71 % (s.d = 0.63, ****p-value < 0.0001) respectively, corresponding to ∼10.95-fold and 13.2-fold higher editing efficiency compared to that of control replicates. Significance was calculated using one-way ANOVA followed by t-test with Šídák’s multiple comparisons test. b, representation of pegRNA spacers with different edit-to-nick distances. PegRNAs were designed with different edit-to-nick distances (denoted in +). Nick sites indicated within the highlighted boxes. c, bar graph depicting the mean A-to-G on-target correction for each pegRNA with different edit-to-nick distances (+1, +2 and +14). On average PE2-SpRY-Cas9 with (+1) pegRNA exhibited comparatively higher editing efficiency of 21.17 % (s.d = 4.73) and PE2-SpG-Cas9 exhibited 6.14 % (s.d = 2.43, ***p-value < 0.0001). Whereas the combination of (+2) pegRNA and PE2-SpRY-Cas9 yielded 16.54 % (s.d = 1.71) and PE2-SpG-Cas9 7.5 % (s.d = 1.68, ****p-value < 0.0001). In addition, when paired with (+14) pegRNA PE2-SpRY-Cas9 and PE2-SpG-Cas9 yielded 7.57 % (s.d = 0.82) and 4.36 % (s.d = 0.82, p-value = 0.1784) efficiencies respectively. Significance was calculated using two-way ANOVA followed by t-test with Šídák’s multiple comparisons test. d, A-to-G on-target correction when PE2-SpRY-Cas9 is paired with (+1) epegRNA compared to (+1) pegRNA. (+1) epegRNA yielded the highest level of editing efficiency of 27.36 % (s.d = 5.57, p-value = 0.2188) compared to (+1) pegRNA (s.d = 4.73, 21.17 %). Error bars represent the mean with s.d of n = 3 independent biological replicates.

### Proximity of the desired edit to the nick

Additionally, we assessed how the proximity of the desired edit to the nick site influences editing efficiency. We designed pegRNAs with varying edit-to-nick distances (denoted as +1, +2 and +14) (**Figure 1b**) and co-transfected them with either PE2-SpG-Cas9, PE2-SpRY-Cas9 or PE2-SpCas9 (control). Interestingly, the combination of PE2-SpRY-Cas9 and (+1) pegRNA outperformed (21.17 %, s.d = 4.73 and ∼3.45 times more efficient than PE2-SpG-Cas9, p-value < 0.0001) the editing level of PE2-SpG-Cas9 (6.14 %, s.d = 2.43). However, when PE2-SpRY-Cas9 was paired with (+2) pegRNA, we observed a decline in the editing efficiency (16.54 %, s.d = 1.71) compared to that of (+1) pegRNA (21.17% vs 16.54%). In contrast, PE2-SpG-Cas9 with (+2) pegRNA resulted in a comparatively higher editing level than (+1) pegRNA (7.5 %, s.d = 1.68). Further assessment of editing efficiency by pegRNA with longer edit-to-nick distances (+14 pegRNA) revealed reduced editing efficiencies of 7.57 % (s.d = 0.82) and 4.36 % (s.d = 0.82, p-value = 0.1784) for PE2-SpRY-Cas9 and PE2-SpG-Cas9 respectively. Compared to (+1) pegRNA, this is a 2.8 (PE2-SpRY-Cas9) and 1.4-fold (PE2-SpG-Cas9) reduction in editing efficiencies (**Figure 1c**). Overall, these data confirmed that combination of pegRNA with different edit-to-nick distances can manipulate mammalian cell prime editing efficiency and in particular, we observed that PE2-SpRY-Cas9 efficiency tended to increase as the edit-to-nick distance of the pegRNA is decreased.

Next, we tested whether the PE2-SpRY-Cas9 editing efficiency can be further improved by pairing it with (+1) epegRNA. Compared to PE2-SpRY-Cas9 and (+1) pegRNA (control) (21.17%, s.d = 4.7) combination of (+1) epegRNA yielded the most efficient level of editing thus far (27.36 %, s.d = 5.57, p-value = 0.2188 by unpaired t-test) corresponding to a 1.29-fold increase in editing efficiency (**Figure 1d**).

### PegRNA with same sense mutations

To further evaluate the utility of prime editing we paired PE2-SpRY-Cas9 with pegRNAs with same sense mutations (SSM),where an additional silent point mutation is introduced in the reverse-transcription template (RTT).^19^ Here, two sense-pegRNAs were designed by introducing SSM at positions 2 (2SSM) and 5 (5SSM) in the RTT and co-transfected with PE2-SpRY-Cas9. Compared to that of PE2-SpRY-Cas9 and (+1) epegRNA (control, s.d = 5.57, 27.36 %), 2SSM pegRNA resulted in a mean on-target editing of 9.09 % (s.d = 1.46, p-value = 0.0357) whereas combination of PE2-SpRY-Cas9 with 5SSM1 pegRNA resulted in an average editing of 10 % (s.d = 2.18, p-value = 0.1082) (**Figure 2a**). Overall, both 2SSM and 5SSM1 displayed substantially lower editing efficiencies compared to (+1) epegRNA control replicates (**Figure 2b**). These results suggest that incorporation of pegRNA with same-sense mutations in some cases can hinder prime editing efficiency.

**Figure 2.**
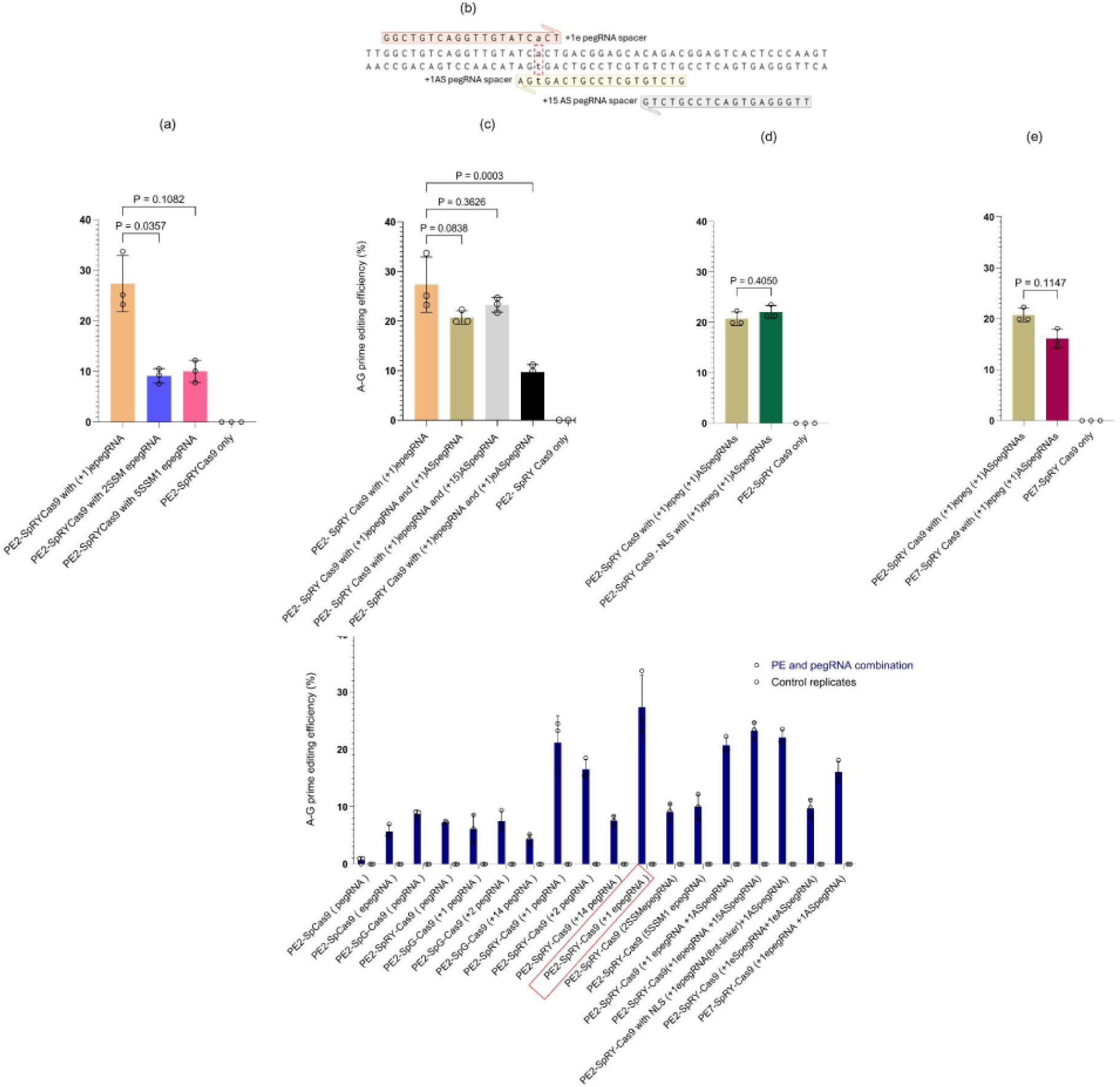
**a**. Bar graph showing in vitro editing efficiencies of PE2-SpRY-Cas9 combined with pegRNAs with varied same sense mutations compared to that of PE2-SpRY-Cas9 and (+1) epegRNA combination (control). Among the two SSM designs tested, mean prime editing efficiencies ranged from 9.09 – 10 % and were, however, lower than that of the control (s.d = 5.57, 27.36 %). Combination of PE2-SpRY-Cas9 with 2SSM pegRNA resulted in a mean on-target editing of 9.09 % (s.d = 1.46,***p-value = 0.0357) whereas combination of PE2-SpRY-Cas9 with 5SSM1 pegRNA resulted in an average editing of 10 % (s.d = 2.18,***p-value = 0.1082). Significance was calculated using one-way ANOVA followed by t-test with Šídák’s multiple comparisons test. **b**. schematic of eHOPE approach representing pegRNAs with varied PAM-in distances at the marine TPP1 p.R207X target site. eHOPE uses a pair of sense (+1) epegRNA and antisense pegRNAs (+1) ASpegRNA and (+15) ASpegRNA encoding the same edit to target both sense and antisense DNA strands respectively. **c**.bar graph depicting the mean A- to-G on-target correction for eHOPE approach compared against PE2-SpRY-Cas9 and (+1) epegRNA combination (control, s.d =5.57, 27.36 %). eHOPE combination of (+1) epegRNA and (+1) ASpegRNA resulted in 20.71 % editing (s.d = 1.34, p-value = 0.0838) whereas (+1) epegRNA and (+15) ASpegRNA resulted in 23.26 % (s.d = 1.47, p-value = 0.3626). Among the three eHOPE pegRNA combinations tested, combination of (+1) epegRNA and (+1) eASpegRNA resulted in relatively lower editing efficiency of 9.73 % (s.d = 1.52, **p-value = 0.0003). Significance was calculated using ordinary one-way ANOVA followed by t-test with Šídák’s multiple comparisons test. **d.** bar graph depicting the comparison of in vitro A-to-G editing efficiencies using PE2-SpRY-Cas9-NLS combined with (+1) epegRNA and (+1) ASpegRNA and compared to editing of PE2-SpRY-Cas9 without the NLS (control, s.d = 1.34, 20.71 %). Combination of PE2-SpRY-Cas9-NLS and (+1) epegRNA yielded a mean on-target editing efficiency of 22.07 % (s.d = 1.25, p-value = 0.4050) corresponding to a 1.07-fold increase in editing than of the control. Significance was calculated using unpaired t-test. **e,** bar graph showing mean A-to-G on-target correction using PE7-SpRY-Cas9 and compared to editing of PE2-SpRY-Cas9 (control, s.d = 1.34, 20.71 %). PE7-SpRY-Cas9, (+1) epegRNA and (+1) ASpegRNA combination resulted in 16.09 % of editing (s.d = 1.83, p-value = 0.01147) corresponding to a 1.29-fold reduction in editing efficiency compared control replicates. Significance was calculated using unpaired t-test. **f.** bar graph shows summary of on-target editing efficiencies resulting from various combinations of PE variants and pegRNAs, compared against controls (transduced only with PE expressing plasmid) and measured by high-throughput sequencing. Error bars represent the meaning with s.d of n = 3 independent biological replicates.

### Homologous 3’ extension mediated prime editor (HOPE)

Next, we utilized a modified homologous 3′ extension mediated prime editing (eHOPE) approach which utilises paired pegRNAs (engineered sense and antisense pegRNAs) encoding the same edits in both sense and antisense DNA strands. Utility of both of these sense (SpegRNA) and antisense (ASpegRNA) pegRNAs permits simultaneous editing of the target loci. ^20^ We designed three different ASpegRNAs with varying PAM-in distances (+1 and +15) and (+1) eASpegRNA (with 3’ motif) and transfected along with (+1) engineered pegRNA (eSpegRNA) and the PE (**Figure 2b**). Editing efficiencies were compared against the control replicates transfected with PE2-SpRY-Cas9 with (+1) epegRNA (27.36 %, s.d = 5.57). On average, combination of (+1)epeg with the (+1) ASpegRNAs yielded an editing level of 20.71 % (s.d = 1.34, p-value = 0.0838), (+15) ASpegRNA resulted in 23.26 % (s.d = 1.47, p-value = 0.3626) and (+1) eASpegRNA resulted in relatively lower editing efficiency of 9.73 % (s.d = 1.52, p-value = 0.0003) (**Figure 2c**). Together these results demonstrate that the eHOPE approach did not contribute to further improving the prime editing efficiency and rather, we observed a marked decline (∼2.8-fold decrease compared to control) in editing when both pegRNAs were engineered variants.

### PE2 with Nuclear localisation signals

Alongside pegRNA structural modifications, we also incorporated strategies aimed at enhancing PE performance. As such, the PE2-SpCas9 architecture was modified with a c-Myc nuclear localisation signal (NLS) sequence.^26^ Robust entry of genome editing agents into the cellular nucleus is critical for editing efficiency and therefore, inclusion of NLS enhance the likelihood of successful genome editing. Compared to PE2-SpRY-Cas9 with (+1) epeg and (+1) ASpegRNAs (control, 20.71 %, s.d = 1.34), PE2-SpRY-Cas9-NLS with (+1) epeg and (+1) ASpegRNAs yielded 22.07% of prime editing efficiency (s.d = 1.25, p-value = 0.4050). Overall, addition of NLS did not improve the prime editing outcome beyond what has already been achieved with PE2-SpRY-Cas9 and (+1) epegRNA combination (27.36 %) (**Figure 2d**).

### La protein

In addition to 3’ trimmed evopreQ1 (tevopreQ1), a small RNA-binding exonuclease protection factor ‘La’ was recently found to promote prime editing efficiency in PE7.^15^ The La protein functionally interacts with the 3′ ends of polyuridylated pegRNAs and thereby prevents degradation of the pegRNA 3’ end.^15^ Guided by this study, we cloned PE7 into SpRY-Cas9 to achieve an improved PAM flexibility and also to reduce the edit-to-nick by 1-3 bases. When PE7-SpRY-Cas9 was co-transfected with (+1) epegRNA and (+1) ASpegRNA a mean editing efficiency of 16.09 % (s.d = 1.83, p-value = 0.1147) was achieved compared to controls treated with PE2-SpRY-Cas9, (+1) epegRNA and (+1) ASpegRNA (20.71 %, s.d = 1.34) (**Figure 2e**). Overall, these results indicate that incorporating PE7 strategy into SpRY-Cas9 did not confer any advantage over PE2-SpRY-Cas9 under the tested conditions.

### Overall PE efficiency

Overall, these results demonstrate that equivalent murine mutation (p.R207X) of the pathogenic human *TPP1* mutation of c.622 C>T (p. R208X) is amenable to CRISPR-prime editing. This study also demonstrates that structural improvements in prime editor components have the potential to improve prime editing in cultured cells. Although no editing was detected with PE2-SpCas9, replacing the Cas9 module with the SpRY-Cas9 variant led to a modest improvement in prime editing efficiency (no editing vs 7.23 % respectively). To further improve the prime editing levels we progressively introduced more potent strategies including modifications to the PE protein ^22,23,26^ and pegRNA structure ^15,17,19,20^ and achieved up to 27.36 % of maximum on-target prime editing (PE2-SpRY-Cas9 with (+1) engineered pegRNA) (**Figure 2f**).

### Indel formation

Undesired insertions or deletions (indels) are a critical drawback of prime editing, as they may induce amino acid changes in the edited sequence. In this study, indel formation was not observed with most of the PE strategies, however, with an exception of pegRNA with same sense mutations (5SSM) (**Figure 3 a, b**).

**Figure 3.**
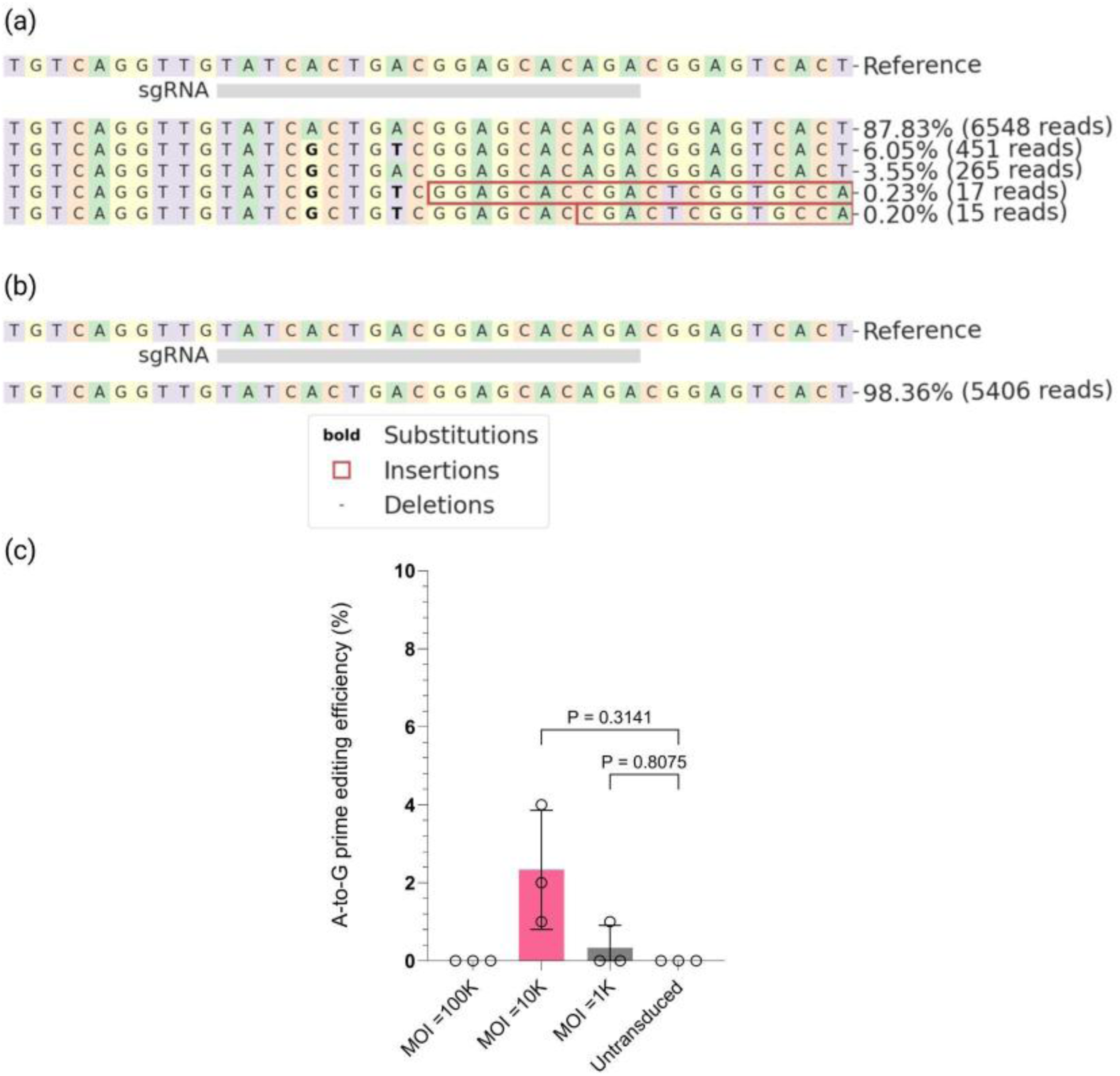
**a, b** Example allele frequency tables generated by CRISPResso2 showing indel formation for PE2-SpRY-Cas9 combined with 5SSM pegRNA (a) compared against the control replicates (b) (transduced only with PE expressing plasmid. Substitutions are shown in bold font, and the insertions are shown within the red box. Nucleotides are indicated by different colours (A=green, C=red, G=yellow, T=purple). **c**. Bar graph showing PE2-SpRY-Cas9-eVLP mediated mean A-to-G on-target correction at different MOIs (100K, 10K and 1K) compared against the non-transduced control replicates. Compared to the control replicates, the highest editing of 2.33 % (s.d = 1.52, p-value = 0.3141) was observed with the MOI=10k, whereas MOI=1k resulted in 0.33 % editing (s.d = 0.57, p-value = 0.8075). Significance was calculated using one-way ANOVA followed by t-test with Šídák’s multiple comparisons test. Error bars represent the mean with S.D of n = 3 independent biological replicates.

### Packaging of PE

#### PE-eVLPs

Although plasmid transfection was effective for initial *in vitro* screening, *in vivo* application of prime editors requires safe and efficient delivery of prime editor and pegRNAs to target cells. Therefore, to realise the maximum therapeutic benefit, the choice of an appropriate delivery vector is critical. Although various delivery systems are currently available, packaging of the prime editor and pegRNA is often restricted due to their large size (7 kb) ^27^. However, recent studies have shown the utility of viral vectors,^28^ lipid nanoparticles^29^ and engineered virus-like particles (eVLPs)^27^ for PE and pegRNA packaging. Among these delivery vectors, engineered VLPs are emerging as a promising delivery vehicle that offer several advantages over viral and non-viral delivery systems. A recent study demonstrated a strategy for progressive optimisation of PE-eVLPs to maximise editing efficiency.^27^ In this study we first attempted simple PE2-SpRY-Cas9 introduction to synthesize PE2-SpRY-Cas9-eVLPs. In cultured HEK293A cells, transfection of PE2-SpRY-Cas9-eVLP packaging plasmids mediated prime editing up to 12.66 % (s.d = 0.57). To further evaluate the editing potential of PE-eVLP platform, we generated PE-eVLPs carrying an engineered sense pegRNA (+1epegRNA) and an antisense pegRNA (+1ASpegRNA) and transduced cultured HEK293A cells harboring the *TPP1* R207X mutation). Cells were transduced at different multiplicity of infection (MOIs) of 1,000, 10,000 and 100, 000 (titer of 2.963 × 10¹¹ TU/mL) and evaluated editing efficiencies. No statistically significant editing compared to controls was detected at any MOI. Given the low level of *in vitro* editing, PE-eVLPs did not qualify for *in vivo* characterisation (**Figure 3c**).

Overall, our findings establish the proof of concept that CRISPR-based prime editing can correct the pathogenic human *TPP1* nonsense mutation of c.622 C>T (p. R208X) associated with CLN2 disease.

## DISCUSSION

Prime editing offers several advantages over base editing and has the greatest versatility for correcting a vast majority of disease-associated mutations that are inaccessible to base editing. PE has already progressed to the clinic, with multiple clinical trials currently underway.^30–34^ In this study, we demonstrated that the equivalent murine mutation (p.R207X) of the pathogenic human *TPP1* nonsense mutation of c.622 C>T (p. R208X) is amenable to prime editing techniques.

Prime editing involves three precise, nucleic acid hybridisation events: the initial pegRNA-target sequence hybridisation, hybridisation of the primer binding site (PBS) and reverse transcription. Priming and high fidelity during these steps are critical to prevent bystander editing.^9,35^ Despite recent advancements, prime editors have thus far not outperformed base editing efficiencies. Typically, PE efficiencies are optimised through structural modifications of its components, including the catalytically impaired Cas9 nickase domain, reverse transcriptase (RT) domain and the pegRNA.^15,17,19,23,26^ This study progressively introduced structural improvements to both PE and pegRNA which resulted in substantially improved prime editing efficiencies, up to 27.36 % *in vitro*.

It was further demonstrated that the choice of Cas9 variant significantly influenced the prime editing efficiencies. During initial screening, engineered SpCas9 variants such as SpG and SpRY outperformed the conventional SpCas9 (0.66 %) achieving up to 8.7 % and 7.2 % of editing efficiencies respectively.

Incorporation of a trimmed prequeosine_1_-1 riboswitch aptamer (tevopreQ_1_) to the 3′ motif indeed improved the prime editing efficiency, and with (+1) epegRNA yielding up to 27.36 %, corresponding to a 1.29-fold increase in prime editing compared to an unmodified (+1) pegRNA (21.17 %). As an alternative to tevopreQ_1_, a frameshifting pseudoknot from Moloney murine leukemia virus (MMLV) has also been reported to prevent 3’ degradation.^17,36^

Additionally, we also observed that the proximity of the desired edit to the nick is critical in determining the editing efficiency. Further attempts were made to improve the PE efficiency via optimized eHOPE method using pegRNA pairs encoding the same edits in the sense and antisense DNA strands.^20^ While all of our eHOPE pegRNA pairs mediated a robust PE efficiency, the combination of epegRNA and (+15) ASpegRNA (with a +15 edit-to-nick distance) resulted in a higher mean editing efficacy (23.26 % of A-to-G editing) across the 3 combinations.

It was also demonstrated that the addition of N-terminal c-Myc nuclear localisation signals at the C terminus of the PE protein did not improve the editing efficiency (efficiency 22.07 % with NLS vs. 20.71 % without NLS, p-value = 0.4050).

To achieve the maximal therapeutic potential *in vivo*, PE will need to be paired with a suitable delivery system. Much progress has been achieved utilising adeno-associated viral (AAV) vectors for *in vivo* base editing. For example, AAV base editing strategies have been used to attenuate disease pathologies in mice models of human genetic diseases such as metabolic liver disease, Duchenne muscular dystrophy, and spinal muscular atrophy.^37–40^ Despite their promising therapeutic potential, packaging of the PE into a single AAV is challenging owing to its’ large size .^28^,^41^

To bypass the packaging limitations of AAV vectors, PE components can be delivered via eVLPs. PE-eVLPs have been structurally optimised to package dual pegRNAs, cargo release, and localisation. In particular, the optimised v3 and v3b PE-eVLPs have previously been shown robust editing of pathogenic mutations in the murine brain ^27^. *In vitro* results from this project demonstrated that PE-eVLPs (simple replacement of base editors (BEs) with PEs in the optimised BE-eVLP system) did not achieve editing above the background in controls.^27^

While we achieved proof-of-concept correction of the R207X mutation, several promising PE variants remain to be tested. PE4 and PE5 strategies ^13^, which incorporate mismatch repair modulation, and the more compact PE6d,^14^ optimised for dual AAV delivery, could offer further improvements in both efficiency and therapeutic applicability. Future experiments could be done to assess these newer variants in combination with dual AAV delivery systems.

Overall, the data establish that equivalent murine mutation (p.R207X) of the pathogenic human *TPP1* nonsense mutation of c.622 C>T (p. R208X) is amenable to prime editing technique. By systematically introducing structural changes to the PE and the pegRNA correction of the mutation was possible, with an editing efficiency of up to 27.36 % *in vitro*. It is anticipated that the correct combination of a delivery system will warrant pre-clinical evaluation and potentially contribute to the development of prime editing as a genetic treatment for Ceroid neuronal lipofuscinosis type 2 (CLN2).

## METHODS

### Generation of stable cell line

To facilitate screening of the gene editing agents a stable cell line was generated by introducing the equivalent murine mutation (p.R207X) of the pathogenic human *TPP1* nonsense mutation of NM_000391.4 (TPP1):c.622C>T; p.Arg208Ter into the HEK293A genome using a lentiviral vector. First, we curated and identified the sequence of human *TPP1* gene and aligned it with mouse ortholog.

### Generation of lentivirus

To generate lentivirus, HEK293FT producer cells were plated (R70007, ThermoFisher Scientific) at a density of 6 x 10 ^ 6 /dish and supplemented with Dulbecco’s modified Eagle medium fortified with high glucose (Life Technologies), 10 % (vol / vol) Fetal Bovine Serum (Life Technologies), 1 % (vol / vol) antibiotic-Penicillin-Streptomycin (Thermofisher Scientific) and incubated and maintained in a humidified chamber at 37°C with 5 % CO_2_. At 70 % confluence, HEK293FT cells were co-transfected with 10 µg of p-delta 8.91 lentiviral packaging plasmid, 1 µg of pDMG envelope plasmid and 10 µg of transfer plasmid using 63 µl of TransIT 2020. Transfection complexes were added to the cells dropwise and following a 6-hour incubation at 37°C with 5 % CO_2_, media was replaced with fresh DMEM supplemented with 10 % FBS. At 48 hours post-transfection, lentivirus-containing supernatant was collected, filtered through a Millex-HV 0.45-μm low protein-binding membrane (Millipore) and concentrated using pegIT (PEG-it™ Virus Precipitation Solution (5×) Cat. # LV810A-1/ LV825A-1).

### LV transduction of HEK293A cells, selection and validation

HEK293A cells were seeded in a 24-well culture plate at a density of 40k/well incubated at 37°C with 5 % CO_2_. At 30-40 % confluence HEK293A cells were transduced with a premade mixture of lentivirus and lentiblast premium (1:200 dilution) at the multiplicity of infection (MOI) of 0.3. After 48 hours of transduction, existing media was replaced with DMEM containing 1 µg/mL puromycin. Both transduced and non-transduced cells (CONTROL) were checked daily to monitor the degree of cell death within non-transduced wells.

When CONTROL cells had completely died, surviving cells were expanded onto a larger vessel (T75 flask) and Sanger verified (forward primer GAACAAAAACTCATCTCAGAAGAGGATCTG and the reverse primer AGCGTAATCTGGAACATCGTATGGGTA). Once the successful integration of the p.R207X mutation sequence was confirmed, cell aliquots were cryopreserved for future experiments.

### Molecular cloning

A major determinant of prime editing efficiency is the design of the pegRNA. A web tool, PrimeDesign (https://drugthatgene.pinellolab.partners.org/)^42^ was used to design the equivalent murine mutation (p.R207X) of the pathogenic human *TPP1* nonsense mutation of NM_000391.4 (TPP1):c.622C>T; p.Arg208Ter.

Additionally, when designing the various pegRNA variants with structural modifications, parameters such as the strand orientation, editing position relative to the CRISPR/Cas nicking site (+1, +2, +14, +15; positive denotes the upstream of the target edit), length of the primer binding site (PBS), reverse transcriptase template length (RTT), sense (S) and antisense (AS) spacer and extension sequences were provided to generate potential pegRNA sequence candidates.

To further improve the prime editing efficiency we designed 3 different sense-pegRNAS (spegRNA) with same-sense mutations (SSM) introduced at different positions in the RT-template as described by Li *et al*, 2022.^19^ Oligonucleotide sequences for extension sequences, spacer, scaffold and 3’ pseudoknot were annealed and assembled with digested pU6-pegRNA-GG-acceptor (Addgene, #132777) vector backbone.

Each pegRNA contains a spacer, scaffold, RTT, PBS and 3’structural motif elements and is under the control of a U6 promoter. Engineered pegRNAs were designed by incorporating a trimmed prequeosine1-1 riboswitch aptamer (tevopreQ1) to the 3′ terminus as described by Nelson *et al*, 2022.^17^ For homology 3’ extension-mediated prime editing (HOPE) paired pegRNAs encoding the same edits in both sense and antisense DNA strands were designed.^20^ Isothermal assembly of these pegRNA elements is described below.

Briefly, vector backbone pU6-pegRNA-GG-acceptor (Addgene, #132777) was used for pegRNA and epegRNA cloning. 2000 ng of pU6-pegRNA-GG-acceptor plasmid (addgene Plasmid #132777) was digested at 37 °C for 16-17 hours, using 1µl of BsaI -HFv2 (NEB), 3 µl of 10 x Cutsmart buffer, and H_2_O according to the manufacturer’s protocol. Digested ∼2.2kb fragment was gel purified by gel electrophoresis with a 1 % agarose gel using Monarch® DNA Gel Extraction Kit (NEB #T1020.^9^ In separate reactions 1µL of 100 µM of spacer, caccGAGCCCACATCTTTGGCTGTCgtttt (spacer oligo top), ctctaaaacGACAGCCAAAGATGTGGGCTC (spacer oligo bottom) scaffold, and PBS/RTT oligonucleotides were annealed using 23µL of annealing buffer (H2O supplemented with 10 mM Tris-Cl pH 8.5 and 50 mM NaCl), then heated at 95 ^0^C for 3 minutes and cooled gradually (0.1 ^0^C/s) to 22 ^0^C.

Annealed oligonucleotides were diluted to 1:4 with 75 µL H_2_O to a final concentration of 1 µM and proceeded for golden gate assembly.^9,10^ 1 µL of gel purified vector backbone (at 30 ng/µL), 1 µL of spacers (1 µM), pegRNA 3’extension (gtgcGCTCCGTCAGcGATACAACCTGACAGCCAAAGATGTGGG (Extension oligo top), aaaaCCCACATCTTTGGCTGTCAGGTTGTATCgCTGACGGAGC (Extension oligo bottom)), and scaffold were assembled using 0.25 µL of BsaI-HFv2 (NEB), 0.50 µL T4 DNA ligase (NEB), 1 µL of 10x T4 DNA ligase buffer (NEB) and H_2_O. The reaction mix was incubated for 5 min at 16 ^0^C and 5 min at 37 ^0^C for 8 cycles in a thermocycler and followed by 15 min at 37 ^0^C, then 15 min at 80 ^0^C, then held at 12 ^0^C.^9,10^

### Cell culture

All cell culture was performed in 37°C 5 % CO_2_ controlled incubators and in tissue culture hoods, passaging cells no more than 20 times. Cells were cultured in Dulbecco’s modified Eagle medium (DMEM) with high glucose (Life Technologies) and transfected at a density of 80,000 cells per well in a 12-well tissue culture-treated plate (In Vitro Technologies). All the plasmids for mammalian cell transfections were prepared using PureLink™ HiPure plasmid midiprep kit (Catalogue K210005).

Cells were transfected using TransIT 2020 (Catalog number selected: MIR 5404). Cells were co-transfected with 750 ng of prime editor and a 250 ng of sgRNA or pegRNA (125 ng of sense and 125 ng of antisense pegRNA) plasmids and cells were harvested 96 hours post-transfection unless otherwise stated. Genomic DNA was isolated using Zymo Quick-DNA Miniprep (Catalogue# D3024) according to the manufacturer’s information. Supernatant was aspirated and 500 µl of genomic lysis buffer was added directly to the cell monolayer. Cell pellets were resuspended by vortexing 4-6 seconds and letting stand for 5-10 minutes at room temperature. Samples were incubated at room temperature for 5-10 minutes and the samples were transferred to a Zymo-Spin™ IICR column in a collection tube. Sample mixture was centrifuged at 10,000 x g for one minute and the flow-through was discarded. Next, the spin column was transferred to a clean microcentrifuge tube a 200 µl of DNA pre-wash buffer was added to the spin column and centrifuged at 10,000 x g for one minute. 500 µl of g-DNA wash buffer was then added to the spin column and centrifuge at 10,000 x g for one minute. Lastly, the spin column was transferred to a clean microcentrifuge tube and 30 µl DNA of the elution buffer was added to the spin column. Followed by an incubation for 2-5 minutes at room temperature DNA was eluted by centrifuging 30 s at top speed.

### Plasmid transfections

Cells were plated in 48-well plates (Corning,3548) at a density of 50 000 cells per well. Following the manufacturer’s protocol, cells were transfected using Mirus TransIT® 2020 (Catalog number: MIR 5404). Cells were co-transfected with 750 ng of prime editor plasmids and 250 ng of *CLN2* mutation targeting pegRNA plasmid (s). Genomic DNA samples were amplified using the following amplification primers, (TCGTCGGCAGCGTCAGATGTGTATAAGAGACAGGAACAAAAACTCATCTCAGA AGAGGATCTG and GTCTCGTGGGCTCGGAGATGTGTATAAGAGACAGAGCGTAATCTGGAACATCGTATGGGTA including 5’ extensions to enable Illumina barcoding primer amplification).

12.5 μl of Q5® High-Fidelity DNA Polymerase, 1.25 μl of 10 μM primers, 100 ng of genomic DNA as the template and H_2_O up to 25 μl was added per PCR reaction. Cycling conditions were 98 °C for 30 s, then 30 cycles of (98 °C for 10 s, 62 °C for 30 s, and 72 °C for 20 s) followed by a final extension of 1 min at 72 °C.

### HTS of genomic DNA samples and data analysis

Cells were harvested 96 hours post-transfection unless otherwise stated. Genomic DNA was isolated using Zymo Quick-DNA Miniprep (Catalogue# D3024). Genomic DNA samples were amplified using amplification primers including 5’ extensions to enable Illumina barcoding primer amplification. 12.5 μl of Q5® High-Fidelity DNA Polymerase, 1.25 μl of 10 μM primers, 100 ng of genomic DNA as the template, and H2O up to 25 μl was added per PCR reaction.

Cycling conditions were 98 °C for 30 s, then 30 cycles of (98 °C for 10 s, 62 °C for 30 s, and 72 °C for 20 s) followed by a final extension of 1 min at 72 °C.

To evaluate the base editing efficiencies, purified PCR samples were further proceeded with either Miseq sequencing or Sanger sequencing using BigDye Terminator v3.1 Cycle Sequencing kit (Cat No. 4337454, Life Technologies). Briefly, the reaction was set as follows: 1.75 μl 5× Sequencing Buffer, 0.25 μl BigDye® Terminator v3.1 Ready Reaction Mix, 1 μl of 10 μM diluted primer 1 ng/μL DNA template, and PCR-grade water to a final reaction volume of 10 μl and sequenced using the protocol: 1 min at 95°C (30 s at 95°C, 30 s at 56°C, and 1 min at 60°C) × 25 cycles, and hold at 15°C. Sequencing reactions were analysed on an Applied Biosystems 3500 DNA Analyzer. Whereas for Miseq sequencing, samples were barcoded with i5 and i7 indices using Nextera XT Index Kit v2 set A-D (Illumina). 2 μl of PCR template, 2.5 μl of i5 and i7 adapters, 12.5 μl of Q5 Hot Start High-fidelity 2× master mix, and 5.5 μl of PCR-grade water for a final volume of 25 μl was prepared. Thermocycling conditions were programmed to an initial denaturation at 95 °C for 2 min, 15 cycles of cycling denaturation at 95 °C for 15 s, 61 °C annealing for 20 s, and 72 °C extension for 20 s, followed by 72 °C final extension for 2 min. Followed by 1× paramagnetic bead cleanup step samples were quantified using either Qubit™ dsDNA BR Assay Kit (Life Technologies) or Qubit™ 1X dsDNA HS Assay Kit (Cat No. Q33230, Thermofisher scientific). Purified samples were then normalized to 4 nM based on the calculation of the amplicon size. 5 µL of each library are pooled into a final library that is validated using High Sensitivity D1000 ScreenTape (Agilent Technologies). According to the manufacturer’s instructions, 4 nM of the pooled library was diluted to 12 pM, 10 % PhiX control was added and finally, samples were loaded onto a Miseq V3 flow cell (Paired-end; 2 × 300, 600 cycles) (Cat No. MS-102-3003, Illumina). Sequencing outputs were then joined and analysed using the CRISPResso2 (V.2.0.29) workflow.^43^ The following quantification window parameters were used: -w 20 -wc -10. Base editing efficiencies were reported as the percentage of sequencing reads containing a given base conversion at the specific positions within the editing window. Sanger sequencing of PCRs from genomic DNA was analysed using EditR.^44^

## STATISTICAL ANALYSIS

Data are reported as mean ± s.d. unless stated otherwise. Significance was calculated using two-way ANOVA followed by t-test with Šídák’s multiple comparisons test. The number of biological replicates and statistical tests are described in the figure legends. All statistical tests were performed using GraphPad Prism 10 https://www.graphpad.com/.

## ACKNOWLEDGEMENT

AWH is supported by an Australian National Health and Medical Research Council Leadership Fellowship. We are grateful for funding from Retina Australia and the Batten Disease Support and Research Association Australia.

## AUTHOR CONTRIBUTIONS

All authors contributed equally.

